# Valence-specific EEG microstate modulations during self-generated affective states

**DOI:** 10.1101/2023.09.23.559103

**Authors:** Karina Nazare, Miralena I. Tomescu

## Abstract

We spend a significant part of our lives navigating emotionally charged mind-wandering states by spontaneously imagining the past or the future, which predicts general well-being. We investigated brain self-generated affective states using EEG microstate analysis to identify the temporal dynamics of underlying brain networks that sustain endogenous affective state activity. With this aim, we compared the temporal dynamics of five distinct microstates between baseline resting-state, positive (e.g., awe, contentment), and negative (e.g., anger, fear) affective self-generated states. We found affect-related modulations of B, C, and D dynamics. Microstates B and D were increased, while microstate C was decreased during negative and positive valence self-generated affective states. In addition, we found valence-specific mechanisms of spontaneous affective regulation. Negative valence self-generated affective states specifically modulate the increased presence of D microstates and decreased occurrence of E microstates compared to baseline and positive valence affective states. The self-generated positive valence affective states are characterized by more prevalent B and les present A microstates compared to both baseline and negative valence affective states. These findings provide valuable insights into the neurodynamic patterns of affective regulation and implications for developing biomarkers for therapeutic interventions in mood and anxiety disorders.

## Introduction

Emotional or affective states play an essential role in our well-being by modulating our thoughts, behaviors, and social interactions. Moreover, we spend a significant part of our daily life navigating spontaneously self-generate affective states by imagining past or future scenarios to adapt to challenging life contexts or enjoy rewarding experiences impacting decision-making and general well-being (Andrews-Hanna et al., 2013; Killingsworth & Gilbert, 2010; Ruby et al., 2013). Many studies focused on emotions elicited by external stimulations without personal context (e.g., audio, visual, audio-visual stimulation). However, substantial inter-individual variability in the neural, physiological, and behavioral results challenges these methods based on imposed categorical boundaries between emotions (Barrett, 2006; Lindquist et al., 2012). To overcome these limitations and accommodate the inter-individual variability from the subjective experience versus the normative categorization of stimuli, we can focus on self-generated affective states based on naturalistic depictions of real-life emotion-induction context scenarios (Wilson-Mendenhall et al., 2013).

Only a few studies investigated self-generated affective states. Using a method closer to the spontaneous mind-wandering affective elicitation, Onton and Makeing (2009) developed a study where participants self-induced several affective states following verbally guided narrative suggestions (Onton & Makeig, 2009). Using spatiotemporal decomposition of the EEG, authors reached small to high accuracy in emotional classification based on independent-component analysis (ICA) decomposition of EEG signal (Hsu et al., 2022; Kothe et al., 2013). However, the complex multidimensional ICA discriminated between self-generate affective states when compared to relaxation and only at the individual level (Hsu et al., 2022; Kothe et al., 2013). During the affective states, ICs in prefrontal, sensorimotor, premotor, and higher-level visual brain areas differentiated between affective and relaxed states (Hsu et al., 2022; Kothe et al., 2013). However, no difference was noted between negative and positive affective states despite clear differentiations in the fMRI literature pointing towards valence-specific modulations where the dorsolateral prefrontal cortex (DLPFC), frontal pole, rostro-dorsal anterior cingulate cortex (ACC), and supplementary motor area predicted negative valence states and regions of the reward circuit such as the midbrain, ventral striatum, and caudate nucleus activated more during the positive valence states (Colibazzi et al., 2010).

An alternative method to investigate valence-specific brain spatiotemporal dynamics during affective states, and, at the same time, taking advantage of the high temporal resolution and more naturalistic approach of the EEG, is by employing the EEG microstates analysis to the available dataset from Onton and Making (2009). EEG microstate reveals the fast-changing temporal dynamics of resting state networks with high temporal resolution (Lehmann et al., 1987; Michel & Koenig, 2018). EEG microstate dynamics discriminate between different cognitive states like mental calculation, visualization, verbalization, and autobiographical memory, and socio-affective states and traits (Bréchet et al., 2019; Milz et al., 2016; Schiller et al., 2023; Seitzman et al., 2017; Tarailis et al., 2023). With a high degree of reproducibility, four (A-D) and seven EEG microstates (A-G) have been identified across many conditions and participants. Accumulating evidence suggests that EEG microstates represent the electrical fingerprints of resting-state networks; however, their one-to-one correspondence is still debated (Britz et al., 2010; Custo et al., 2017; Michel & Koenig, 2018; Musso et al., 2010; Yuan et al., 2012). Generally, A-B microstates are related to bottom-up visual and auditory/language-related. Microstates C, D, and E have been associated with core regions of top-down functional networks such as the default mode (DMN), the dorsal attention (DAN), and salience networks (SN) (Michel & Koenig, 2018). Cognitive and socio-affective manipulation manipulations significantly mediate DMN, DAN, and SN-associated microstates (Schiller et al., 2023; Tarailis et al., 2023).

Few studies investigated how temporal dynamics of EEG microstates advance our understanding of affective processing and regulation. With the goal of decoding emotional states, temporal structures of microstates were used to classify between arousal and valence with approximately 65% accuracy (Chen et al., 2021). Other studies show that C microstate coverage and the occurrence of microstate B were essential for recognizing discrete positive and negative emotions (Liu et al., 2023; Shen et al., 2020). Another study showed that microstate D displayed a negative association with valence (Shen et al., 2020). In a recent study utilizing a video-watching paradigm, the temporal dynamics of C and D microstates dissociate high versus low valence and arousal (Hu et al., 2023). While the C microstate showed a positive relation with arousal, microstate D occurred more often during negative valence videos (Hu et al., 2023).

Different methodological approaches might explain the contradictory results. For example, the high inter-individual variability of emotional reactivity to external stimuli might account for the lack of consistency. To overcome this challenge, we can focus on the neural mechanisms of self-generated affective states that naturally occur during our daily spontaneous mind-wandering. Moreover, self-generated affective state modulations based on naturalistic life scenarios might reveal unique microstate dynamics patterns reflecting spontaneous endogenous affective state regulation. With this aim, we investigated spatiotemporal microstate changes between baseline resting-state, positive (awe, compassion, contentment), and negative (anger, disgust, fear) affective self-generated states. Disentangling these brain dynamics might be essential in understanding basic affective and mood state regulation patterns that predict well-being.

## Methods

### Dataset Experimental Paradigm

In this study, we analyzed an EEG dataset collected and described in previous research by Onton and Makeig (Onton & Makeig, 2022). The study employed guided imagery to facilitate participants’ self-induction of several affective states. High-density EEG data was collected during the elicitation and maintenance of these affective states (Onton & Makeig, 2009).

At the beginning and end of the session, participants were instructed to rest for two minutes, summing up to four minutes of non-affective baseline recording. Then, participants received instructions on the button press to self-report their affective states, thus marking the beginning and the end of each affective state before transitioning to the next state. Participants had to interoceptively detect and pay attention to salient somatosensory sensations to heighten and extend the duration of their emotions. Finally, participants listened to voice recordings that advised visualizing scenarios they had previously encountered or would experience to self-induce the affective states. There were seven negative valences, undesirable target emotions (anger, disgust, fear, frustration, grief, jealousy, and sadness), and eight positive valences, pleasant target emotions (awe, compassion, contentment, excitement, happiness, joy, love, and relief), presented in an alternate pseudo-randomized order (Onton, 2009; Onton & Makeig, 2022).

### Dataset Acquisition

The EEG data were acquired from 250 scalp, four infraocular, and two electrocardiographic electrodes at a sampling rate of 256 Hz using the Biosemi AtiveTwo EEG system. Individual locations of the electrode positions in 3D (x, y, z) coordinates were provided for each participant. For more details, see Onton and Makeig 2009 (Onton & Makeig, 2009). Participants sat in a comfortable chair in a dimly-light, quiet room, received audio instruction, and listened to the narrative descriptions using earbuds. Participants were instructed to extend the duration of the affective state naturally. This resulted in experimental sessions that lasted roughly 80 minutes. The 480 recorded affective states (15 emotions x 32 subjects) ranged from 43 seconds to 12 minutes (on average, 218 ± 94 seconds) (Onton & Makeig, 2009, 2022).

### Dataset Participants

Thirty-four participants (14 male, 19 female; age range: 18–38 years; age mean and standard deviation: 25.5 ± 5 years) volunteered for this experiment at the University of California, San Diego (UCSD). All participants gave informed consent, and the experimental procedures complied with the institutional requirements of UCSD. Given the self-paced, self-induced nature of the paradigm, authors report that all participants could experience affective states for most of the studied emotions by using either their self-generating strategies and imagination or following the voice narrative recording suggestion. For more details, see Onton and Making 2009 (Onton & Makeig, 2009).

In the present article paradigm, the evoked affective state was considered felt only after the corresponding button press. As no button press makers were identified for subject 33, he has been excluded from further analysis.

### Data processing

The dataset included preprocessed data with excluded noisy channels of electrodes with poor skin contact and a 1-Hz high pass filter. We first continued pre-processing the data by applying a 40 Hz low-pass Butterworth filter. Next, the affective state epochs were exported for each subject using the marked beginning and end of each affective experience. These were further ordered and concatenated into three categories: baseline, positive affective, and negative affective states.

The preprocessed data were subjected to the Infomax independent component analysis (ICA) (Jung et al., 2000), and based on visual inspection of the topography and amplitude of the components, we eliminated cardiac and eye movement artifacts. ICA was carried out on the entire dataset, including data from the marginal electrodes, allowing for more precise identification of eye, pulse, and muscular tension components.

After excluding noisy channels in the original database preprocessing, the retained electrode numbers varied between 134 and 235. To unify the space across subjects while reducing the computational load of further investigations, we reduced the number of electrodes to the standard 128 channels (Biosemi). First, the ECG and ocular electrodes were identified and eliminated from the analysis. Then, we interpolated the EEG tracks from the individual electrode 3D space maps by selecting the front, top, back, left, and right landmarks (H22, B12, C14, A31, D20) to the 128 Biosemi 3D common standard coordinate system for statistical analysis across subjects (FpZ, Cz, Oz, T7, and T8). The Cartool interpolation tool, using a 3D spline interpolation that accounts for the actual geometry of the head, was used for interpolation (Brunet et al., 2011). Finally, before microstate analysis, the data were downsampled to 128 Hz and re-referenced to the average reference for further analysis.

### Microstate analysis

Microstate analysis mainly consists of two stages: first, the clustering of EEG data to find the most representative template maps, which correspond to the different microstates, and second, fitting them back to the EEG data to quantify their temporal parameters.

The EEG topographies surrounding the local maxima of the Global Field Power (GFP) exhibit the highest signal-to-noise ratio (Murray et al., 2008). Topographies corresponding to GFP peaks were submitted to a modified k-means cluster analysis to identify the most representative classes of stable topographies. GFP represents the global pattern of brain activity and is defined as follows:

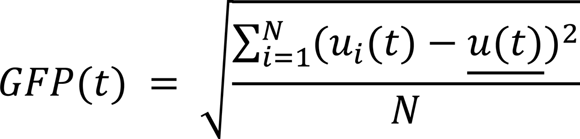

where *N* is the number of channels, *ui*(*t*) is the measured amplitude of a channel at a time *t*, <u>*ui*(*t*)</u> is the average voltage of the *N* electrodes at the specific time *t* (Murray et al., 2008).

The K-means clustering was carried out in two stages, first at the individual level and then at the group level, by clustering each individual dominant topography. The ideal number of clusters at the individual and affective conditions was determined using the Cartool, based on seven maximally independent criteria: Davies and Bouldin, Gamma, Silhouette, Dunn Robust, PointBiserial, Krzanowski-Lai Index, and Cross-Validation (Brunet et al., 2011; Custo et al., 2017).

Then, each subject’s recorded brain electrical activity is modeled as a time sequence of microstates. After identifying microstate topographies for each condition (**Fig. 1**), during the fitting process of the microstates, the entire EEG of the participants was used, excluding only the marked artifact epochs. A temporal smoother with the following settings was applied: Besag factor of 10, window half-size 3 (24 ms), and rejection of brief time intervals (of less than three timeframes or 23.4 milliseconds). Each time point of the individual data was assigned to the microstate cluster with which it correlated best to measure the temporal parameters of microstates. Short periods of noise in the data were eliminated using a 0.7 correlation coefficient threshold. The analysis excluded these periods (Brunet et al., 2011). For each affective state condition (positive or negative) and baseline, we estimated the occurrence and duration of each microstate in each individual. The average uninterrupted time (in ms) that a specific microstate map was present, or the time the subject remained in that state, is called *mean duration (ms)*. A microstate’s occurrence (Hz), independent of its mean duration, reveals how frequently a specific microstate occurs every second.

**Figure 1.**
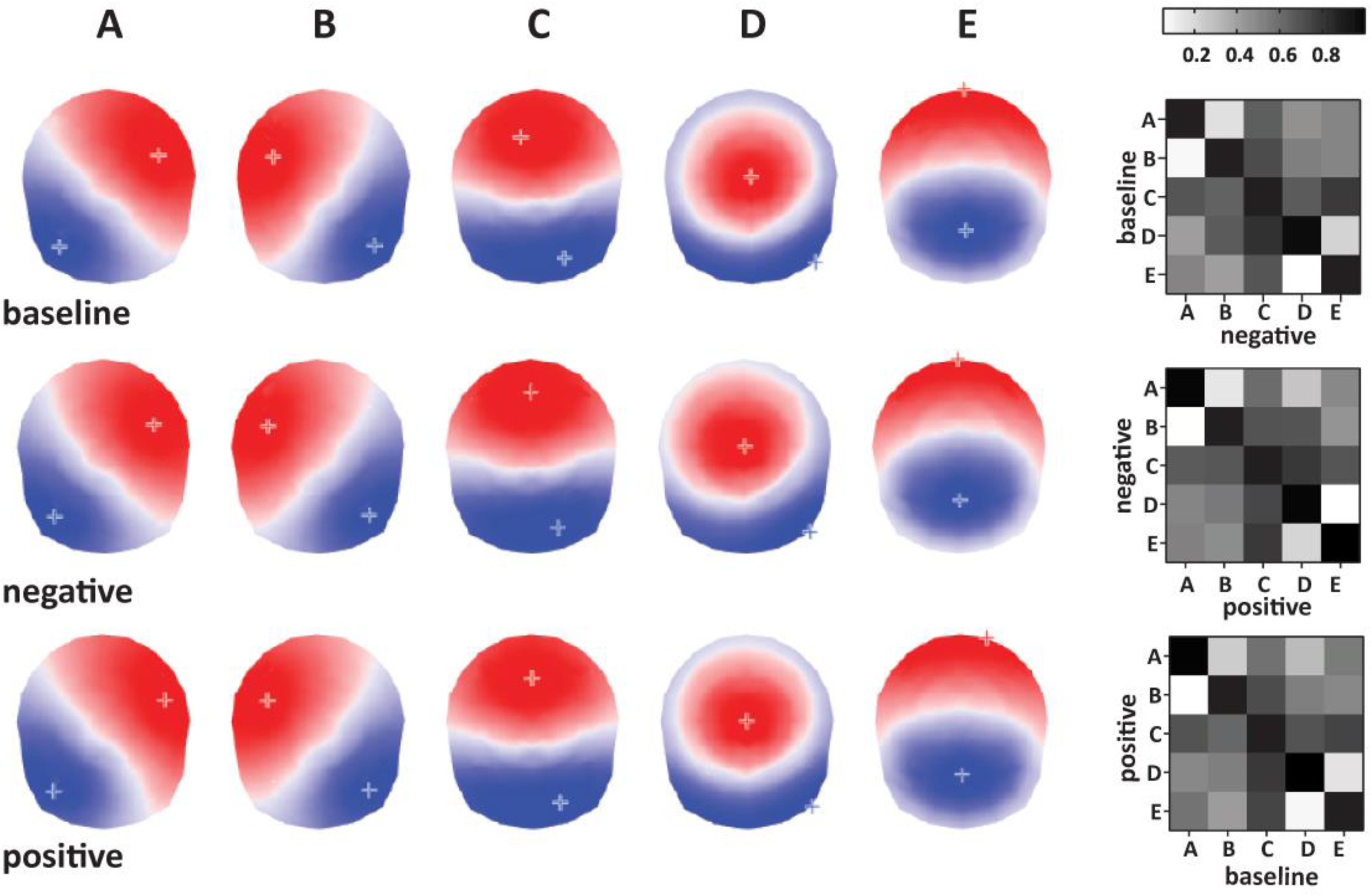
Left: Microstate topographies identified for each condition (top to bottom: baseline, negative affect, and positive affect conditions). Right: Microstate correlation coefficient matrixes used for labeling and microstate ordering between conditions.

A thorough truncation method was used to establish comparability between the data produced from the affect categories and the baseline, in which the signals during negative and positive affects corresponded precisely with the briefest baseline period. As a result of this methodological approach, only the first 31350 timeframes (4.09 min) of the recording, presenting only the first alphabetical emotions of each individual, were maintained for statistical analysis. Specifically, the number of emotions considered for each subject varied based on the duration of the evoked emotion period, with only the first one to three emotions included. This approach encompassed highly arousing negative emotions such as *anger*, *disgust*, and *fear* and positive emotions like *awe*, *compassion*, and *contentment*. One participant (27) did not auto-evaluate himself as attaining the compassion emotion (by not pressing the corresponding button). Thus, we included the next positive emotion, *excitement*.

The free academic software Cartool, Matlab academic software, and EEGlab Matlab plugin were used for the EEG data processing and microstate analysis.

## Results

### Topographic analysis

Using a set of seven independent criteria (detailed in the Method section), we determined that five microstates can optimally describe the group topographical variability (baseline 87.1%, negative affect 87.2%, positive affect 86.6% of explained variance). We labeled them A-E and ordered them according to the between-condition correlation coefficients shown in **Figure 1**.

### Temporal dynamic analysis

The temporal dynamic analysis is based on the temporal properties of each microstate relative to the affective state, including their *occurrence* (Hz) and *mean duration* (ms) (Table 1). To examine how the self-generated affective state (active affective, i.e., negative (NEG), positive (POS), or baseline (BAS) impact temporal properties of microstates, Wilcoxon signed-rank two-tailed tests with FDR correction for multiple comparisons were used.

**Table 1.** Descriptive statistics of *mean duration* (ms) and *occurrence* (Hz) of microstates during baseline, positive and negative affective states.

|  | Microstate | Mean duration (ms) |  | Occurrence (Hz) |  |
| --- | --- | --- | --- | --- | --- |
|  |  | Mean | Standard deviation | Mean | Standard deviation |
| BAS | A | 66.25 | 6.39 | 1.68 | 0.67 |
|  | B | 65.96 | 4.85 | 1.72 | 0.69 |
|  | C | 101.12 | 24.41 | 3.6 | 0.51 |
|  | D | 63.82 | 5.32 | 1.42 | 0.63 |
|  | E | 68.26 | 7.05 | 1.88 | 0.79 |
| NEG | A | 65.96 | 5.95 | 1.73 | 0.71 |
|  | B | 67.34 | 5.74 | 1.77 | 0.68 |
|  | C | 97.11 | 21.30 | 3.72 | 0.56 |
|  | D | 66.49 | 5.89 | 1.76 | 0.63 |
|  | E | 64.71 | 7.19 | 1.68 | 0.82 |
| POS | A | 65.03 | 5.64 | 1.55 | 0.66 |
|  | B | 68.93 | 6.88 | 1.97 | 0.71 |
|  | C | 96.50 | 23.51 | 3.66 | 0.55 |
|  | D | 64.24 | 5.72 | 1.63 | 0.66 |
|  | E | 66.82 | 7.78 | 1.88 | 0.88 |

### Results for microstates *mean duration* (ms)

We found significant differences in the *mean duration* of microstate B, which increased significantly during both POS and NEG affective states vs. baseline (**Fig. 2**, **Table 2**). In addition, the microstate’s B *mean duration* also proved significantly higher during the POS than the NEG. During NEG and POS microstates, the C *mean duration* decreased significantly compared to BAS but did not show significant differences between the NEG and POS conditions. D microstates were significantly longer in NEG when compared to BAS and POS, while E microstates significantly lasted for a shorter amount of time in NEG compared to BAS and POS (**Fig. 2**, **Table 2**).

**Figure 2.**
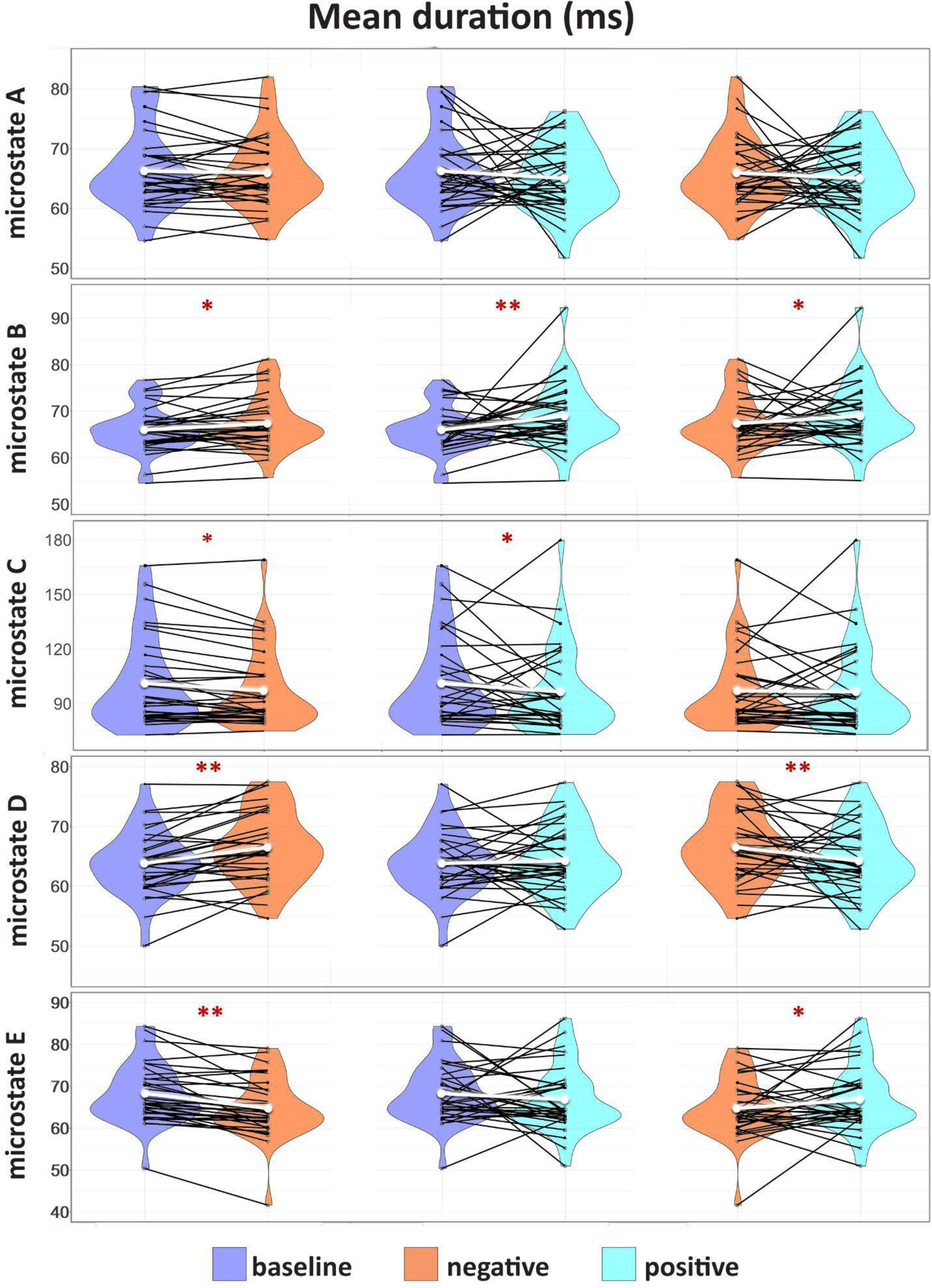
Results for microstates *mean duration* (ms). Group comparison of the distribution and the median of the *mean duration* (ms) of each identified microstate fitted for BAS, NEG, and POS. Intercategorically connected black points represent intra-individual data; the median is represented in white. The significant differences are marked with *p<0.05, ** p<0.001.

**Table 2.** Results for microstates *mean duration* (ms)

|  | Microstate | Wilcoxon - T | Wilcoxon - Z | Confidence Interval | p-value (FDR corrected) | Cohen's *d* |
| --- | --- | --- | --- | --- | --- | --- |
| BAS - NEG | A | 247 | 0.59 | [0.05 to 0.54] | 0.54 | 0.04 |
|  | B | 151 | 2.31 | [0.02 to 0.04] | **0.02** | **0.25** |
|  | C | 138 | 2.54 | [0.01 to 0.03] | **0.01** | **0.17** |
|  | D | 89 | 3.42 | [0.00 to 0.02] | **0.001** | **0.47** |
|  | E | 41 | 4.27 | [0.00 to 0.01] | **0.00009** | **0.49** |
| BAS - POS | A | 170 | 1.97 | [0.04 to 0.03] | **0.08** | **0.20** |
|  | B | 69 | 3.77 | [0.00 to 0.01] | **0.0007** | **0.49** |
|  | C | 105 | 3.13 | [0.00 to 0.02] | **0.004** | **0.19** |
|  | D | 233 | 0.84 | [0.05 to 0.39] | 0.39 | 0.07 |
|  | E | 204 | 1.36 | [0.04 to 0.17] | 0.21 | 0.19 |
| NEG - POS | A | 185 | 1.7 | [0.04 to 0.08] | 0.1 | 0.15 |
|  | B | 149 | 2.34 | [0.01 to 0.03] | **0.03** | **0.25** |
|  | C | 243 | 0.67 | [0.05 to 0.50] | 0.5 | 0.02 |
|  | D | 59 | 3.97 | [0.00 to 0.01] | **0.0003** | **0.38** |
|  | E | 100 | 3.22 | [0.00 to 0.02] | **0.0031** | **0.28** |

### Results for microstates *occurrence* (Hz)

Microstate A occurs much less frequently in POS than both BAS and NEG, while the microstate B occurrence rate is significantly higher in POS than in BAS or NEG. There are no significant differences in microstate C occurrence rates for either category. In the case of microstate D, the occurrence rate is significantly higher in POS compared to BAS, NEG compared to BAS, and NEG compared to POS. Microstate E is significantly lower in NEG than both BAS and POS. (**Fig. 3**) The statistical results of the temporal dynamic analysis are presented in detail in **Table 3**.

**Figure 3.**
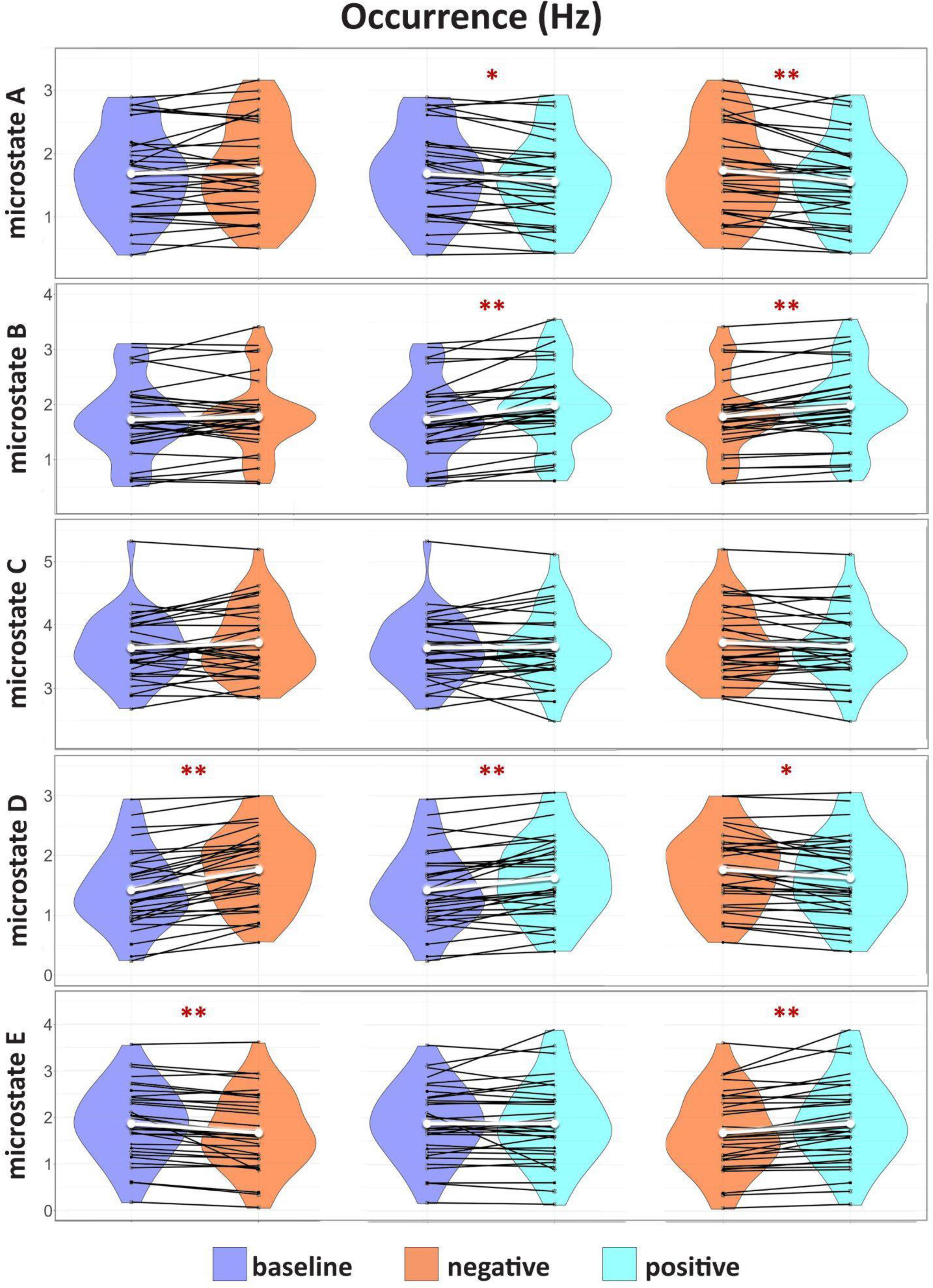
Results for microstate *occurrence* (Hz). Group comparison of the distribution and the median of the *occurrence* (Hz) of each identified microstate fitted for BAS, NEG, and POS. Intercategorically connected black points represent intra-individual data; the median is represented in white. The significant differences are marked with *p<0.05, ** p<0.001.

**Table 3.** Results for microstate *occurrence* (Hz)

|  | Microstate | T | Z | Confidence interval | p-value<br>(corrected) | Cohen'<br>s d |
| --- | --- | --- | --- | --- | --- | --- |
| BAS -<br>NEG | A | 242 | 0.68 | [0.05to 0.49] | 0.49 | 0.07 |
|  | B | 215 | 1.17 | [0.04 to 0.24] | 0.3 | 0.08 |
|  | C | 196 | 1.5 | [0.03 to 0.13] | 0.21 | 0.15 |
|  | D | 20 | 4.65 | [0.0 to 0.01] | **0.00001** | **0.53** |
|  | E | 28 | 4.51 | [0.00 to 0.02] | **0.00001** | **0.24** |
| BAS -<br>POS | A | 113 | 2.99 | [0.00 to 0.03] | **0.004** | **0.19** |
|  | B | 22 | 4.61 | [0.00 to 0.01] | **0.00001** | **0.36** |
|  | C | 237 | 0.77 | [0.04to 0.43] | 0.54 | 0.11 |
|  | D | 61 | 3.92 | [0.00 to 0.02] | **0.0002** | **0.31** |
|  | E | 278 | 0.04 | [0.05to 0.96] | 0.96 | 0.002 |
| NEG -<br>POS | A | 79 | 3.6 | [0.00 to 0.03] | **0.0005** | **0.26** |
|  | B | 36 | 4.36 | [0.00 to 0.01] | **0.00006** | **0.28** |
|  | C | 214 | 1.18 | [0.05to 0.23] | 0.23 | 0.04 |
|  | D | 115 | 2.95 | [0.00 to 0.04] | **0.003** | **0.20** |
|  | E | 64 | 3.86 | [0.00 to 0.02] | **0.0002** | **0.23** |

## Discussion

Self-generated affective states significantly change the resting pattern of spontaneous microstates with small to medium effect sizes that reflect general affect and valence-specific mechanisms of spontaneous affective regulation. Self-generated affective states lead to modulations of microstates B, C, and D dynamics. Microstates B and D showed increased presence during negative and positive valence self-generated affective states. We also found a decreased presence of C microstates during affective states compared to baseline resting state. The results align with brain sources of microstates C, pointing towards association with task-negative posterior DMN core regions like the posterior cingulate cortex, precuneus, and left angular gyrus (Custo et al., 2017). Based on the functional relevance of microstate C and previous findings of its temporal reduction during states of cognitive or behavioral manipulations (Tarailis et al., 2023), the reduction of C microstates during self-generated affective states might reflect a general re-organization pattern of mind-wandering and a possible more aroused, goal-oriented pattern of thought. Microstate C increases during states of relaxation and correlates with more profound states of mind-wandering (less thought discontinuity, less verbal thought about self) after a Social Imitation task (Tomescu et al., 2022). Indeed, we previously found a negative association between C microstates and self-oriented, verbal, and discontinuous patterns of thoughts associated with decreased self-reported levels of stress (Tomescu et al., 2022). In addition, Pan et al. (2020) also found a negative association between microstate C and rumination level. The authors argue that microstate C might be a reliable index of ruminations (Pan et al., 2020). Ruminations are negative valence impulsive patterns of thoughts and an essential mind-wandering transdiagnostic factor in all mood disorders (McLaughlin & Nolen-Hoeksema, 2011; Pan et al., 2020). In our data, we observed a decreased C microstates *mean duration* for positive and negative valence affective states, suggesting an association with more aroused resting states compared to baseline. Additionally, our results align with Hu et al., showing decreased microstate C after emotional audio-visual tasks without a main effect of valence (Hu et al., 2023). In parallel to a decreased presence of C microstates, they also found an increased presence of B and D microstates (Hu et al., 2023). We confirm their findings, showing that during sef-generated positive and negative valence affective states, there is an increase in the presence of B and D microstates. Moreover, we extend their findings, showing a decreased presence of A and E microstates and valence-specific microstates modulations.

The valence-specific modulation points towards a mechanism by which self-generated positive valence affective states are characterized by more prevalent B and les present A microstates compared to baseline and negative valence affective states. Microstate A and B are most often associated with bottom-up networks, auditory-language related and visual activity in the temporal and cortex, left-right cuneus, inferior, and middle occipital gyrus (Britz et al., 2010; Custo et al., 2017; Michel & Koenig, 2018; Tarailis et al., 2023). These microstates tend to increase in presence with more engagement during visualization, verbalization, and autobiographical memory tasks. However, their initial one-to-one functional separation into A verbal-auditory and B visual-related microstates is more complicated than initially reported; see Tarailis et al. (2023) for more details (Tarailis et al., 2023). Regarding microstate A, more convergent literature points towards an association with arousal and alertness (Tarailis et al., 2023). For example, Antonova et al. (2022) found a positive correlation between the mean duration of microstate A and subjective levels of alertness (Antonova et al., 2022). At the same time, other researchers found positive associations between microstate A and prosocial behavior (Schiller et al., 2020). We previously found an increased microstate A presence after both Social Imitation and the control activity of a self-guided arm movement task (Tomescu et al., 2022). Additionally, after the social imitation task, results varied as a function of extraversion (Tomescu et al., 2022). Following this line of associations, compared to baseline or negative valence affective states, the decreased microstate A presence during positive self-generated affective states might also suggest a more relaxed, less alert arousal during a positive affective resting state. Moreover, the microstate A decreased presence results might be specific for the self-generated type of positive affect states as not observed after emotion-inducing videos (Hu et al., 2023). Conversely, microstate B increased during both self-generated positive valence affective experiences and, as previously reported, after emotion-inducing videos (Hu et al., 2023). Additionally, we observed an increased presence of microstate B during negative affective states. As the functional relevance of microstate B has been related to autobiographical memory and scene visualization (Tarailis et al., 2023), the results here might suggest a reflection of the more engagement of these strategies during the self-generated positive affective states. However, significant negative correlations have been found in mood disorder patients with depression scores (Atluri et al., 2018; Yan et al., 2021). Recently, we conducted a meta-analysis performed on clinical studies suggesting the increased B microstate in patients might reflect a compensatory mechanism as larger effect sizes were observed in unmedicated mood disorder patients (Chivu et al., 2023). We suggested that mood and anxiety disorder patients might engage too often in visually related past experiences such as ruminative thought patterns, which fail to compensate for the mood and anxiety symptoms and negatively impact mental health (Chivu et al., 2023). Indeed, microstate B presence was positively associated with self-related thoughts about self-behavior and feelings (Zanesco et al., 2020). However, the results here might suggest that patients could also engage in past experiences of positive valence to compensate for depressive mood.

Negative valence self-generated affective states specifically modulate the increased presence of D microstates and decreased occurrence of E microstates compared to baseline and positive valence affective states. Microstate D is among the four canonical microstates. Studies investigating cognitive state modulations on temporal dynamics of microstates support the view that microstate D is associated with the dorsal attention network involving allocation and maintenance of attentional resources (Britz et al., 2010; Custo et al., 2017; Michel & Koenig, 2018; Tarailis et al., 2023). For example, studies report that microstate D is more present when participants are asked to perform demanding cognitive tasks, such as mental serial subtraction tasks based on focused states of attention (Bréchet et al., 2019; Seitzman et al., 2017). More importantly, microstate D is less present in socially induced spontaneous relaxed states (Tomescu et al., 2022) and shows reduced presence with altered states of attention, consciousness, and lack of cognitive control, such as during auditory-verbal hallucinations in SZ patients (Kindler et al., 2011), deep hypnosis (Katayama et al., 2007), sleep, and dreaming (Bréchet et al., 2020; Brodbeck et al., 2012). Most importantly, the temporal dynamics of microstate D are consistently altered in mood and anxiety patients, showing significant negative associations with depressive symptomatology (Chivu et al., 2023; Murphy et al., 2020). Given our results here, the increased D microstates during negative valence self-generated affective states, support the view that mood disorders might arise from a failure to down-regulate negative emotions. In addition, microstate D quantifiers positively correlate with alertness and reaction time scores in a non-clinical population (Zanesco et al., 2020). Thus, microstate D increase might be related to attention and cognitive control neural resources during emotional attention-demanding tasks. We show that positive and negative valence affective states increased D microstates in line with previous reports of D modulations after emotional-inducing stimuli (Hu et al., 2023). Moreover, based on the association of D microstates with the dorsolateral attention network (DAN), our results are in line with previous ICA-derived prefrontal activation during affective state modulations on the same dataset when compared to relaxed states (Hsu et al., 2022; Kothe et al., 2013). Additionally, we extend these observations by showing that during negative valence affective states, we see a significantly increased presence when compared to the positive affective states, which might reflect the attentional negative valence bias reported in the literature, where negative valence stimuli attract more attentional resources (Baumeister et al., 2001). These results are also in line with fMRI studies on self-generated emotional states showing that unpleasant emotions induced greater activation in a set of regions that included the dorsolateral prefrontal cortex, frontal pole, mid-rostral-dorsal ACC, and supplementary motor area (Colibazzi et al., 2010). These activations might subserve functions like attention allocation, executive functioning, goal-oriented behavior, and emotional regulation during responses to threat-related stimuli (Colibazzi et al., 2010). One study investigated stress-related modulation of EEG microstates and found an increased presence of D microstate and, more significantly, an increased transition between D and salience network-related E microstates (Hu et al., 2023). Moreover, these patterns of increased transitions negatively correlated with salivary cortisol (Hu et al., 2023), further suggesting a possible important role of salience-related and D microstates during negative valence affective states and emotional regulation. Although both microstate C and E microstates were previously associated with salience processing when more than five microstates are represented in the data, microstate E is more associated with the task-positive salience resting-state network with core regions in the superior frontal gyrus, bilateral middle prefrontal cortices ACC and insular cortices (Britz et al., 2010; Custo et al., 2017; Michel & Koenig, 2018; Tarailis et al., 2023). Functionally, microstate E was previously related to the processing of interoceptive and emotional information with increased presence during negative valence effective states, such as after stress exposure, and with increased cognitive load tasks (Hu et al., 2023; Tarailis et al., 2023). Thus, we expected our results to show an increased presence during self-generated affective states. However, these surprising results might be related to the unpredictability nature of the stress exposure and efforts to down-regulate the autonomic system and the nature of the high level of vigilance after stress exposure. Altered E microstates were noted in post-traumatic stress disorder patients, further supporting the association of E microstates with negative valence affect and anxiety (Chivu et al., 2023; Terpou et al., 2022). Terpou et al. (2022) proposed that the brain regions functionally related to the salience network and decreased E microstates might reflect a failure to map relevant bottom-up stimuli, resulting in a hypervigilance state in patients suffering from anxiety-related disorders like PTSD (Terpou et al., 2022). Following the same line of thought, our decreased E microstate during negative valence affective states might be specifically associated with the integrated nature of self-generate negative valence affective states that do not require active salience processing and autonomic activation of the hypothalamic-pituitary-adrenal axis (HPA) for adaptation to stressful contexts. However, more studies are needed to sustain this interpretation and association with successful emotional regulation.

By examining EEG microstates in the context of self-induced affective states, we show valence-specific microstate modulation that extends previous and fast-growing socio-emotional microstate literature. More studies are needed to see how these modulations are influenced by inter-individual emotional regulation traits, clinical symptomatology, and socio-emotional context to sustain general well-being and mental health. However, our findings already provide valuable insights into the neural aspects of emotional regulation and their potential implications for therapeutic interventions in emotional disorders.

## Acknowledgements

We thanks the participants to the study and the authors Onton and Making (2009) for initiating, collecting and making the dataset available (Onton & Makeig, 2009). We extend our gratitude to volunteer students in the lab Octavian F. Mirică and Cosmina A. Duțică, that supported this research during their lab internship and researchers Alexandra Sofonea, and Alina A. Chivu.

## Author’s contributions

M.I.T attracted necessary funding. M.I.T identified the dataset, conceptualized the study and supervised N.K. into conducting the pre-processing and clustering of the microstate analysis, performed the fitting and statistical analyses, wrote the introduction and discussion of the manuscript. N.K. designed and prepared the Figures, wrote the first draft of the methods and results section. All authors contributed to the revision of the manuscript.

## Declarations

Competing interests. The authors have nothing to disclose.

